# New insights into the domain of unknown function DUF of EccC_5_, the pivotal ATPase providing the secretion driving force to the ESX5 secretion system

**DOI:** 10.1101/2024.01.26.577026

**Authors:** Fernando Ceballos-Zúñiga, Margarita Menéndez, Inmaculada Pérez-Dorado

## Abstract

Type VII secretion (T7S) systems, also referred to as ESAT6 secretion (ESX) systems, are molecular machines that have gained great attention due to their implication in cell homeostasis and host pathogen interactions in mycobacteria. The latter include important human pathogens such as *Mycobacterium tuberculosis* (Mtb), the etiological cause of human tuberculosis and a pandemic accounting for more than 1 million deaths every year. The ESX5 system is exclusively found in slow-growing pathogenic mycobacteria, where it mediates the secretion of a large family of virulence factors, the PE and PPE proteins. The secretion driving force is provided by EccC_5_, a multidomain ATPase operating through four globular cytosolic domains, an N-terminal domain of unknown function (EccC^DUF^) and three FtsK/SpoIIIE ATPase domains. Recent structural and functional studies of ESX3 and ESX5 systems have revealed EccC^DUF^ as an ATPase-like fold domain with potential ATPase activity, and whose functionality is essential for secretion. Here we report the crystal structure of *Mtb*EccC_5_^DUF^ domain at 2.05 Å resolution, which unveils a nucleotide-free structure with degenerated *cis*-acting and *trans*-acting elements involved in ATP-binding and hydrolysis. Our crystallographic study, together with a biophysical assessment of *Mtb*EccC_5_^DUF^ interaction with ATP/Mg^2+^, supports the absence of ATPase activity proposed for this domain. We show that this degeneration is also present in DUF domains of other ESX and ESX-like systems, which are likely to exhibit poor or null ATPase activity. Moreover, and based on an *in-silico* model of *Mtb*EccC_5_ N-terminal region, we propose that *Mtb*EccC_5_^DUF^ is a degenerated ATPase domain that may have retained the ability to hexamerise. Observations that call the attention on DUF domains as structural elements with potential implications in the opening and closure of the membrane pore during the secretion process.

## Introduction

Molecular machines are sophisticated protein complexes ubiquitously present in all cellular organisms (Miller & Enemark, 2016). They convert the chemical energy resulting from nucleoside triphosphate (NTP) hydrolysis into the mechanical force needed in numerous cellular events such as DNA replication, protein degradation, cell motility or protein secretion, among others (Miller & Enemark, 2016; Schmidt *et al*., 2012; Famelis *et al*., 2023; Crosskey *et al*., 2020). Members of the superfamily of ATPases Associated with various cellular Activities (AAA +) are key constituents of many molecular machines, where the mechanical work is performed at the expense of adenosine triphosphate (ATP) hydrolysis (Miller & Enemark, 2016; Leipe *et al*., 2002). Type VII secretion systems (T7SSs), also referred to as ESAT6 secretion (ESX) systems, are AAA + dependent molecular machines found in the *Actinomycete* phylum that have gained great attention due to their implication in cell homeostasis and host pathogen interactions in mycobacteria (Famelis *et al*., 2023; Bunduc *et al*., 2020; Houben *et al*., 2014). The latter include devastating pathogenic species such as *Mycobacterium tuberculosis* (*Mtb*), the etiological cause of human tuberculosis (TB) and a pandemic responsible of more than 1 million deaths every year (World Health Organization, 2022). T7SSs are also found in bacteria belonging to the *Firmicutes* phylum (T7SSb), which comprise other important pathogens such as *Staphylococcus aureus*, *Listeria monocytogenes*, *Bacillus anthracis* or *Bacillus subtilis* (Zoltner *et al*., 2016; Mietrach *et al*., 2020).

Mycobacteria produces up to five ESX secretion systems (T7SSa), named from ESX1 to ESX5, consisting of large membrane complexes spanning the inner bacterial membrane. Among them, ESX5 is found almost exclusively in slow-growing pathogenic mycobacteria, where it participates in nutrient uptake, intracellular colonization, and modulation of the immune response during infection through the secretion of a large family of protein effectors, the PE and PPE proteins (Bunduc *et al*., 2020; Houben *et al*., 2014). These roles have important implications in pathogen life cycle and virulence, thus pointing out at the ESX5 as a potential drug target (Bunduc *et al*., 2021). Efforts aiming at the structural-functional characterization of mycobacterial ESX secretion systems, have resulted in the structures of ESX5 from *Mtb* and *Mycobacterium xenopi* (*Mxp*), and ESX3 from *Mycobacterium smegmatis* (*Msm*), which have enlightened the architecture of the pore complex (Famelis *et al*., 2019; Poweleit *et al*., 2019; Bunduc *et al*., 2021; Beckham *et al*., 2021)(Fig. 1A). The latter can be defined as a hexamer of protomers, where each protomer is formed by four different membrane proteins, EccB:EccC:EccD:EccE, interacting with a 1:1:2:1 stoichiometry. Among them, EccC is a pivotal component of the pore complex providing the secretion driving force via ATP hydrolysis (Famelis *et al*., 2019; Poweleit *et al*., 2019; Bunduc *et al*., 2021; Beckham *et al*., 2021). In ESX2-5 systems, EccC consists of two N-terminal transmembrane (TM) helices connected to two additional helices (stalk region) and a domain of unknown function (EccC^DUF^), which are followed by three FtsK/SpoIIIE AAA + ATPase domains, here referred to as D1, D2 and D3 (EccC^D1-3^) (Famelis *et al*., 2023; Bunduc *et al*., 2021) (Fig. 1B). This multi-domain organization is also shared by EccC orthologues (EssC) found in *Firmicutes* (Mietrach *et al*., 2020), which differs in ESX1, where EccC is replaced by two functionally equivalent fragments: EccC_1_a, containing of the DUF and D1 domains; and EccC_1_b containing D2 and D3 domains (Famelis *et al*., 2023; Houben *et al*., 2014). EccC/EssC enzymes are expected to operate through a mechanism involving their hexamerisation. In line with this, the cytosolic region of *Mtb*EccC_5_ has been observed to transient from an extended/open state to a contracted/closed state, where the enzyme multimerises to form the cytosolic chamber that is expected to accommodate the effectors to be secreted (Bunduc *et al*., 2021) (Fig. 1A).

**Figure 1.**
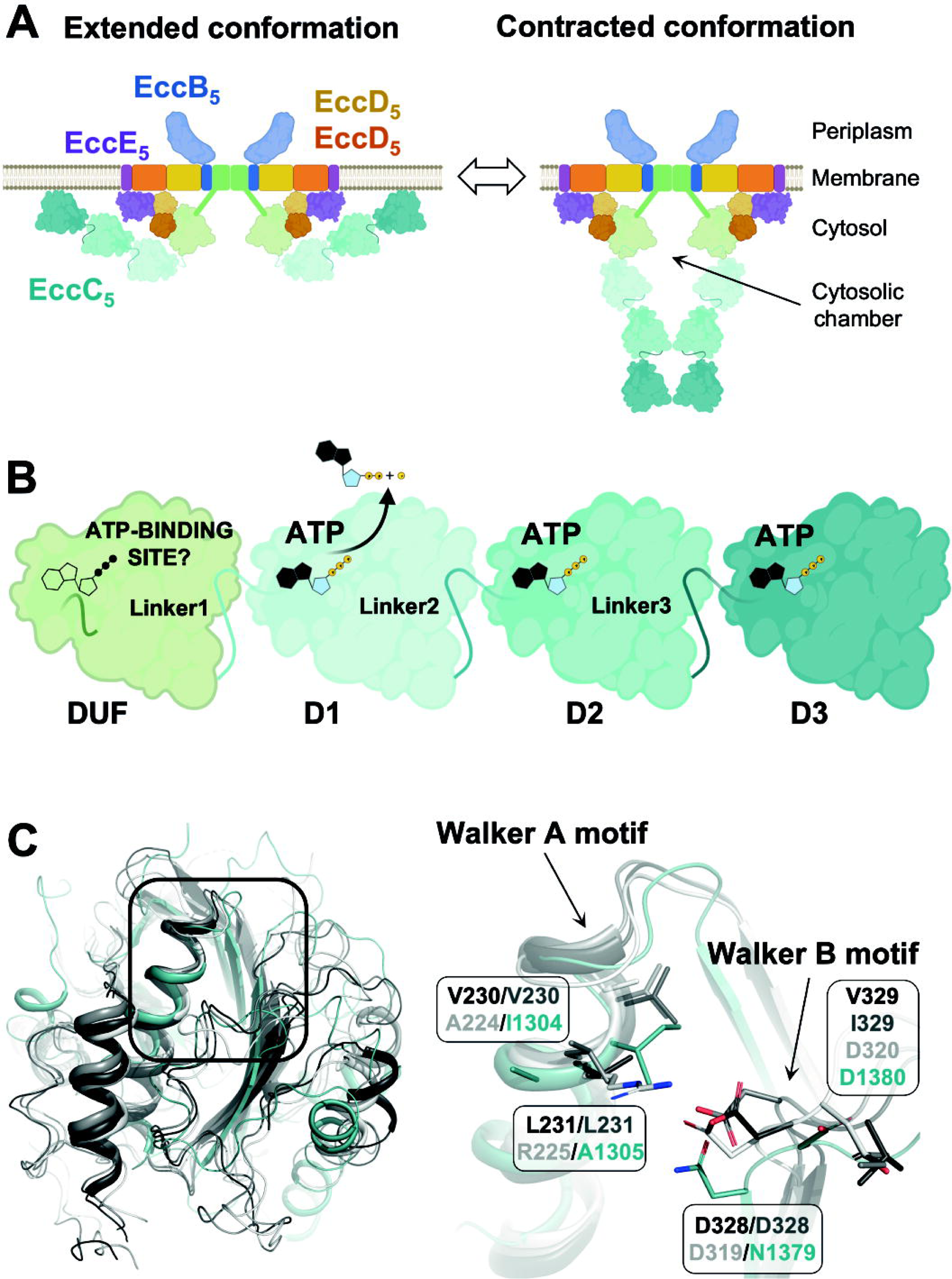
(A) *Mtb*ESX5 model based on cryo-EM reported structures, showing the proposed conformational alteration of EccC_5_ between an open/extended (left) and a close/contracted conformation (right). (B) Architecture of EccC_5_ cytosolic region consisting of the DUF domain followed by three ATPase domains D1, D2 and D3 interconnected by connectors referred to as Linker1-3. (C) Superimposition of *Srs*EssC^D3^ crystal structure (PDB entry 6VT1, in blue) with *Mtb*EccC_5_^DUF^ (PDB entry 7NPR, dark grey), *Mxp*EccC_5_^DUF^ (PDB entry 7B9S, light grey) and *Msm*EccC_3_^DUF^ (PDB entry 6SGX, white) cryo-EM models showing conservation of the overall fold (left) and similar arrangement of both Walker motives (right).

D1-3 domains exhibit a three-layer ɑ-β-ɑ core structure typically found in prokaryotic AAA + enzymes, which is frequently followed by a C-terminal alpha-helical lid domain missing in EccC/EssC enzymes (Zoltner *et al*., 2016). The hexametric architecture observed in ESX3 and ESX5 complexes is in line with the functional oligomeric state frequently found among prokaryotic AAA +. The latter include EccC close relatives, FtsK and SpoIIIE ATPases, reported to function as hexamers, and where ATPase sites locate at the interface between two adjacent protomers (Miller & Enemark, 2016; Leipe *et al*., 2002; Bunduc *et al*., 2021; Rosenberg *et al*., 2015). In such cases, both subunits provide with *cis*- and *trans*-acting catalytic elements required to form the active site. The former includes the Walker A motif, the Walker B motif and Sensor 1, which typically locate at the loop connecting β1 and ɑ1 (P-loop), the C-terminus of the β3 strand, and the loop connecting β4 and ɑ4, respectively. The Walker A motif, also referred to as P-loop, consists of a glycine-rich sequence (‘G_1_xxG_2_xG_3_K[S/T]’ in FtsK homologues) involved in the stabilization of the phosphate group of the nucleotide via highly conserved lysine and serine/threonine residues that enable the correct orientation of the γ-phosphate required for ATP hydrolysis (Miller & Enemark, 2016; Leipe *et al*., 2002). The Walker B motif provides two acidic residues (‘hhhhDE’ in FtsK homologues, where ‘h’ is any hydrophobic amino acid) of which, the second performs as the catalytic base activating the water involved in the nucleophilic attack of the γ-phosphate (Miller & Enemark, 2016; Leipe *et al*., 2002). Sensor 1 is a key catalytic element consisting of a polar residue, typically an asparagine but it can be serine, threonine, or aspartate as well. It locates between both Walker motives to assist in the correct orientation of the catalytic glutamate towards the nucleophilic water. *Trans*-acting elements are positively charged amino acids, mostly (but not exclusively) arginine residues, being thus referred to as arginine fingers or sensors (Miller & Enemark, 2016; Leipe *et al*., 2002). The latter include the Arg finger, which consists of an arginine (or lysine) residue located at the end of the ɑ4 helix that orients towards the neighbouring active site to form contacts with the γ-phosphate. Through this interaction, Arg finger favours the transition state of hydrolysis, thus playing an important role in ATP hydrolysis and, in some cases, in the enzyme multimerization (Miller & Enemark, 2016; Ogura *et al*., 2004). The other two *trans*-acting motifs in AAA + enzymes are Sensors 2 and 3, which are generally arginines involved in sensing and/or stabilising the ATP/ADP-bound state, and promoting conformational changes coupled to these states or nucleotide hydrolysis (Miller & Enemark, 2016; Li *et al*., 2015).

The first structural and functional studies conducted by Rosenberg and co-workers in the EccC from *T. curvata* (TcrEccC) showed that the D1 domain is an active ATPase exhibiting fully conserved Walker A (‘GxxGxGK[S/T]’) and Walker B (‘hhhhDE’) motives with respect to FtsKs, as close canonical orthologues, including the Arg-finger *trans*-acting motif (Rosenberg *et al*., 2015). This and later studies carried out on EccCs/EssCs from Mtb and *S. aureus* showed that Walker A and B motives are, however, degenerated in D2 and D3, which aligns with non-detectable or poor catalytic activities observed for those domains, which are proposed to play a regulatory role (Rosenberg *et al*., 2015; Wang *et al*., 2020; Zoltner *et al*., 2016). More recently, the structural characterization of the *Msm*ESX3 complex revealed that the N-terminal domain of unknown function (DUF) exhibits an ATPase-like fold, also observed in the DUF domains of *Mtb*ESX5 and *Mxp*ESX5 complexes reported afterward (Famelis *et al*., 2019; Bunduc *et al*., 2021; Beckham *et al*., 2021). The authors describe the presence of a Walker B motif (‘hhhhDD’^320^) in *Msm*EccC_3_^DUF^, partially conserved with respect to FtsK orthologs (‘hhhhDE’), and how mutation of each aspartate to alanine obliterates secretion. These observations highlight the essentiality of *Msm*EccC_3_^DUF^ in secretion, and they point to a potential ATPase activity in this domain (Famelis *et al*., 2019). The structure of *Msm*EccC_3_^DUF^ is however devoid of ATP, analogously to *Mtb*EccC_5_^DUF^ and *Mxp*EccC_5_^DUF^ structures that exhibit valine and isoleucine residues replacing the catalytic amino acid, respectively (Famelis *et al*., 2019; Bunduc *et al*., 2021; Beckham *et al*., 2021). These observations expose a variable degeneration of the Walker B motif across different EccC^DUF^ domains that may impair the ATPase activity to a different extent. In addition, *Msm*EccC_3_^DUF^*, Mtb*EccC_5_^DUF^, and *Mxp*EccC_5_^DUF^ domain structures reveal a non-canonical arrangement of the Walker A motif, lacking the typical P-loop structure that is replaced by an extended alpha-helical conformation. Interestingly, this arrangement is observed in the structure of the EssC^D3^ domain of *S. aureus* (*Srs*EssC^D3^, PDB entry 6TV1) also devoid of ATP, and exhibiting an ATP-binding affinity within the low-millimolar range (Mietrach *et al*., 2020).

The structural and functional information available in T7SSs leave outstanding questions regarding the molecular basis underlying the ATPase activity proposed for the DUF domain. In this sense, here we report the crystallographic structure of *Mtb*EccC_5_^DUF^ domain at 2.05 Å resolution, which provides an unambiguous model showing a nucleotide-free structure with degenerated *cis*-acting and *trans*-acting elements involved in ATP-binding and hydrolysis. Our high-resolution structure, together with a biophysical assessment of *Mtb*EccC_5_^DUF^ interaction with ATP/Mg^2+^ *in vitro*, supports the absence of ATPase in this domain. These results are in line with the *in-silico* analysis carried out in other EssC and EccC enzymes, which reveals the presence of degenerated DUF domains in other mycobacterial and non-mycobacterial T7S systems that are likely to exhibit null or deficient ATPase activity. These findings suggest that DUF domains play a different role in the secretion process that, based on an *in-silico* model of *Mtb*EccC_5_ and mutagenesis studies reported in *Msm*ESX3 (Famelis *et al*., 2019), we propose is related to the aperture of the membrane-pore complex during the secretion process.

## Material and methods

### 3.1 Protein expression and purification

Protein constructs containing the DUF domain of EccC_5_ (EccC_5_^DUF^, residues 1-417) from *Mycobacterium tuberculosis* were designed for their recombinant production. Residues 17-118 and 167-198 were replaced by ‘GSSG’ and ‘GSG’ sequences, respectively, in order to remove the transmembrane (TM) and stalk regions and enable its production as a soluble protein. The construct contains a N-terminal 6His-tag followed by a SUMO-tag, and a PreScission 3C cleavage site. The DNA coding for the construct cloned into a pET-28a plasmid (between NcoI and XhoI restriction sites) was purchased to GenScript with codon optimization for *E. coli* expression (pET-EccC_5_^DUF^). Expression of EccC_5_^DUF^ was performed in *E. coli* BL21 (DE3) Star (Invitrogen) using 2xTY media supplemented with kanamycin (50 μg/ml). Cells were grown at 37 °C until an OD_600 nm_ of 0.8 and then cooled down to 16 °C for an hour. Expression was then induced at 16 °C by the addition of 1 mM IPTG for 20 h. Cells were harvested by centrifugation at 5251 *g* for 20 min at 4 °C, and cell pellets were stored at −20 °C.

Cell pellets were thawed and resuspended in 20 mM Tris-HCl pH 7.5 and 500 mM NaCl, 10 mM imidazole (buffer A), supplemented with 5 mM MgCl_2_, 1 mM MnCl_2_ and 1 µg/ml DNAse (Sigma). This step was performed at RT while subsequent purification steps were carried out at 4 °C. Cells were disrupted by sonication and the clarified extract was loaded into a 1 ml HisTrap HP column (Cytiva) previously equilibrated in buffer A. Protein samples were eluted using an imidazole gradient (from 10 mM to 500 mM) and then buffer exchanged to 20 mM Tris-HCl pH 7.5, 150 mM NaCl (Buffer B) using a PD-10 desalting column (Cytiva). Samples were diluted to 1 mg/ml in buffer B prior to overnight cleavage with the 3C protease. The cleaved protein was concentrated using Amicon 10 KDa MWCO concentrators (Millipore), and further purified by size exclusion chromatography using a Superdex 200 (16/60) column (Cytiva) preequilibrated on buffer B or 20 mM HEPES pH 7.5, 150 mM NaCl, 5 mM MgCl_2_ (buffer C), for crystallographic or ITC experiments, respectively. Pure protein fractions obtained from the size-exclusion chromatography (SEC) were pooled, concentrated, and flash-frozen in liquid nitrogen for their storage at −80 °C. Protein sample purity and quantification were assessed using PAGE-SDS and UV-Vis absorbance at 280 nm, respectively.

### 3.2 Protein crystallization, structure resolution and analysis

All crystallization experiments were carried out using the sitting-drop vapour diffusion method at 20 °C in 96-well MRC plates (Hampton Research) and employing an Oryx8 robot (Douglas Instruments). Initial crystals of EccC_5_^DUF^ were obtained in the F2 condition of the Index screen. After optimization, the best crystals grew in 300 nl droplets formed by mixing 150 nl of protein at 10 mg/ml and 150 nl of precipitant condition (25 % (w/v) of PEG MME 2K, 300 mM of trimethylamine N-oxide and Tris-HCl pH 8). Crystals were flash-cooled in liquid nitrogen without cryoprotection for X-ray data collection, which enabled to measure a complete X-ray data set at 100*K* using synchrotron radiation at ALBA (Barcelona, Spain) at 2.05 Å resolution. The data set was indexed and integrated with XDS (Kabsch, 2010), and scaled and reduced using AIMLESS (Evans & Murshudov, 2013). Crystals belonged to the *P*1 space group, with unit cell dimensions *a*=48.30 Å, *b*=52.24 Å, *c*=57.44 Å, ɑ=88.39 °, β=84.60°, γ=80.17 ° (Table 1). Structure resolution was carried out using the molecular replacement method with PHASER (McCoy *et al*., 2007), and the coordinates of EccC_5_^DUF^ structure (1-416 residues) solved by cryo-EM (PDB entry 7NPT) (Bunduc *et al*., 2021) as the search model. A single and unambiguous solution was obtained, which consisted of two EccC_5_^DUF^ molecules in the asymmetric unit. The initial model was subjected to alternate cycles of model building with COOT (Emsley *et al*., 2010), and refinement with PHENIX (Adams *et al*., 2010) and BUSTER (Bricogne G. *et al*., 2017) using NCS restrictions. Electron density maps allowed us to model almost completely both molecules of the asymmetric unit and 287 water molecules. Residues 275-283 and 311-315 (in chain A), and 274-283 (Chain B), were not modelled due to poor electron density observed for these residues. The geometry of the final model was validated using MOLPROBITY(Chen *et al*., 2009). Dihedral angles were analysed with COOT (Emsley *et al*., 2010) and MOGUL (Version 2020.3.0). Figures were generated using PYMOL (Version 2.0; Schrödinger). The atomic coordinates and structure factors have been deposited in the Protein Data Bank under the accession code 8RIN.

**Table 1.** X-ray crystallography statistics.

|  |  |
| --- | --- |
|  | *MtbEccC*<sub>5</sub><sup>DUF</sup> |
| Data collection statistics |  |
| Space group | *P*1 |
| Number of complexes in ASU | 2 |
| *a*, *b*, *c* (Å) | 48.30, 52.24, 57.44 |
| α, β, γ (°) | 88.39, 84.60, 80.17 |
| Wavelength (Å) | 0.9792 |
| Beamline | BL13-XALOC |
| Resolution range (Å) | 57.18 - 2.05 (2.11 - 2.05) |
| *R*<sub>merge</sub> | 0.055 (0.354) |
| *R*<sub>pim</sub> | 0.035 (0.242) |
| Number of total reflections | 114509 (6654) |
| Number of unique reflections | 32869 (2122) |
| Mean (*I*]/*SD*[*I*]) | 9.6 (2.6) |
| *CC*<sub>1/2</sub> | 0.998 (0.869) |
| Completeness (%) | 95.1 (77.9) |
| Multiplicity | 3.5 (3.1) |
| Wilson plot *B* (Å<sup>2</sup>) | 29.5 |
| Refinement |  |
| Number of atoms | 4219 |
| Protein | 3932 |
| Water | 287 |
| *R*<sub>work</sub> (%) | 0.19 |
| *R*<sub>free</sub> (%) | 0.22 |
| <*B*> (Å<sup>2</sup>) | 34.59 |
| rms bond length deviation (Å) | 0.008 |
| rmsd angle deviation (°) | 0.88 |

Values in parentheses correspond to the highest resolution shell.

### 3.3 Differential Scanning Fluorimetry

Differential scanning fluorimetry (DSF) assays were carried out to estimate the inflection temperature (*T*_i_) associated with unfolding transitions of the EccC_5_^DUF^ construct in the absence and in the presence of nucleotide/Mg^2+^. The experiments consisted of monitoring the variation in the emission of fluorescence of tryptophan residues, buried and exposed, at a wavelength of 350 nm and 330 nm, respectively, using the Tycho NT.6 (NanoTemper). DSF experiments were set up to a final volume of 10 µl of buffer C containing the EccC_5_^DUF^ construct at 2 µM in the absence or in the presence of ADP-AlF_3_ at 4 mM. Mixtures of protein with the ATP analogue were incubated for 15 min before DSF measurements. Graph representation and analysis were performed using the Tycho interface of analysis. Three independent runs were used in each case for calculating mean *T*_i_ values and the corresponding standard deviations.

### 3.4 Isothermal Titration Calorimetry

ATP binding and ATPase activity by the EccC_5_^DUF^ domain were assessed by isothermal titration calorimetry (ITC) using an ITC-VP instrument from GE Instruments. EccC_5_^DUF^ at 61 µM was loaded in the cell at 25 °C, and it was titrated with ATP at 1.3 mM (in the syringe) using 15 injections, first and second of 1 and 10 µl and the remaining of 20 µl at intervals of 5 minutes. Both protein and ATP were prepared in buffer C. Analysis of results and final figures were carried out using the Origin 7 ITC Software.

### 3.5 Primary, secondary and tertiary structure analysis *in silico*

A comparative analysis of the putative ATP/Mg^2+^-binding site across DUF domains, and comparison with canonical ATPases, was performed using sequence alignment with Clustal Omega Multi Sequence Alignment (Sievers *et al*., 2011); and by structure superimposition in PYMOL (Version 2.0; Schrödinger) using experimental coordinates deposited in the Protein Data Bank and models predicted by AlphaFold (Jumper *et al*., 2021) available at Uniprot. ATPases used for the sequence alignment correspond to: FtsK from *Pseudomonas aeruginosa* (*Prg*FtsK) (Uniprot entry Q9I0M3; PDB entry 2IUT (Massey *et al*., 2006)); HerA DNA translocase from *Sulfolobus solfataricus* (*Ssl*HerA) (UniProt entry Q97WG8; PDB entry 4D2I (Rzechorzek *et al*., 2014)), MCM helicase from *Pyrococcus furiosus* (*Pfr*MCM) (UniProt entry Q8U3I4; PDB entry 4R7Y (Miller *et al*., 2014)), Vps4 from *Metallosphera sedula* (*Msd*Vps4) (UniProt entry A4YHC5; PDB entry 4D81 (Caillat *et al*., 2015)); EccC from *Thermomonospora curvata* (*Tcr*EccC) (UniProt entry D1A4G7); *Mtb*EccCa_1_ (UniProt entry P9WNB3); *Mtb*EccC_2_ (UniProt entry O05450); *Mtb*EccC_3_ (UniProt entry P9WNA9); *Mtb*EccC_4_ (UniProt entry P9WNA7); *Mtb*EccC_5_ (UniProt entry P9WNA5; PDB entries 7NPT and 7NPR (Bunduc *et al*., 2021)); *Mxp*EccC_5_^DUF^ (UniProt entry I0RZI0; PDB entry 7B9S (Beckham *et al*., 2021)); *Msm*EccCa_1_ (UniProt entry A0A653FCM4), *Msm*EccC_3_ (UniProt entry A0QQ40; PDB entry 6SGX (Famelis *et al*., 2019)); *Msm*EccC_4_ (UniProt entry A0QSN0); EccC from *Geobacillus thermodenitrificans* (*Gth*EccC) (UniProt entry A4IKE7); EssC from *Staphylococcus aureus* (*Srs*EssC) (UniProt entry Q932J9; PDB entry 6TV1); EssC from *Clostridium sp*. (*Csp*EssC) (UniProt entry A0A3R6STG9), EssC *Streptococcus oralis* (*Srl*EssC) (UniProt entry A0A656Z0M6) and YukB from *Bacillus subtilis* (*Bsb*YukB) (UniProt entry C0SPA7). Structural analysis of dihedral angles and their occurrence in the Cambridge Crystallographic Data Centre (CCDC) database was performed using the software MOGUL (Version 2020.3.0).

## 4 Results and discussion

### 4.1 High-resolution structure of EccC_5_^DUF^ domain

The overall crystallographic structure of the *Mtb*EccC_5_^DUF^ domain (Fig. 2A) consists of an ɑ-β-ɑ sandwich highly homolog to the ɑ/β fold found in the core of AAA + domains. The core structure consists of a six-stranded parallel β-sheet (β5-β1-β4-β3-β2-β2’), further extended by an additional two-stranded anti-parallel sheet (β6-β7). This β-sheet core is flanked by two helices (ɑ1 and ɑ2) on one side, and four helices (ɑ2’A, ɑ2’B, ɑ3, and ɑ4) on the other side. The C-terminal ɑ-helical lid present in other prokaryotic AAA + domains is missing in the *Mtb*EccC_5_^DUF^ structure, which is flanked by two parallel helices (ɑA and ɑB) and a two-stranded anti-parallel sheet (βA-βB) that connects with the stalk domain. This structure is conserved in both *Mtb*EccC_5_^DUF^ molecules present in the asymmetric unit, with an rmsd 0.173 Å for 218 Cɑ atoms. Superimposition of crystallographic and cryo-EM (*Mtb*EccC_5_^CRYO^, PDB entry 7NPR (Bunduc *et al*., 2021)) models shows overall structure conservation (rmsd=0.671 Å for 198 Cɑ atoms) except for the N-terminal region (residues 1-16), present but not visible in the crystallographic model, and residues 124-129 and 166-209, connecting to TM and stalk regions, which are truncated in the crystallographic construct (Fig. 2SM). However, despite the overall fold conservation, other differences between our structure and *Mtb*EccC_5_^CRYO^ are found as will be described below.

**Figure 2.**
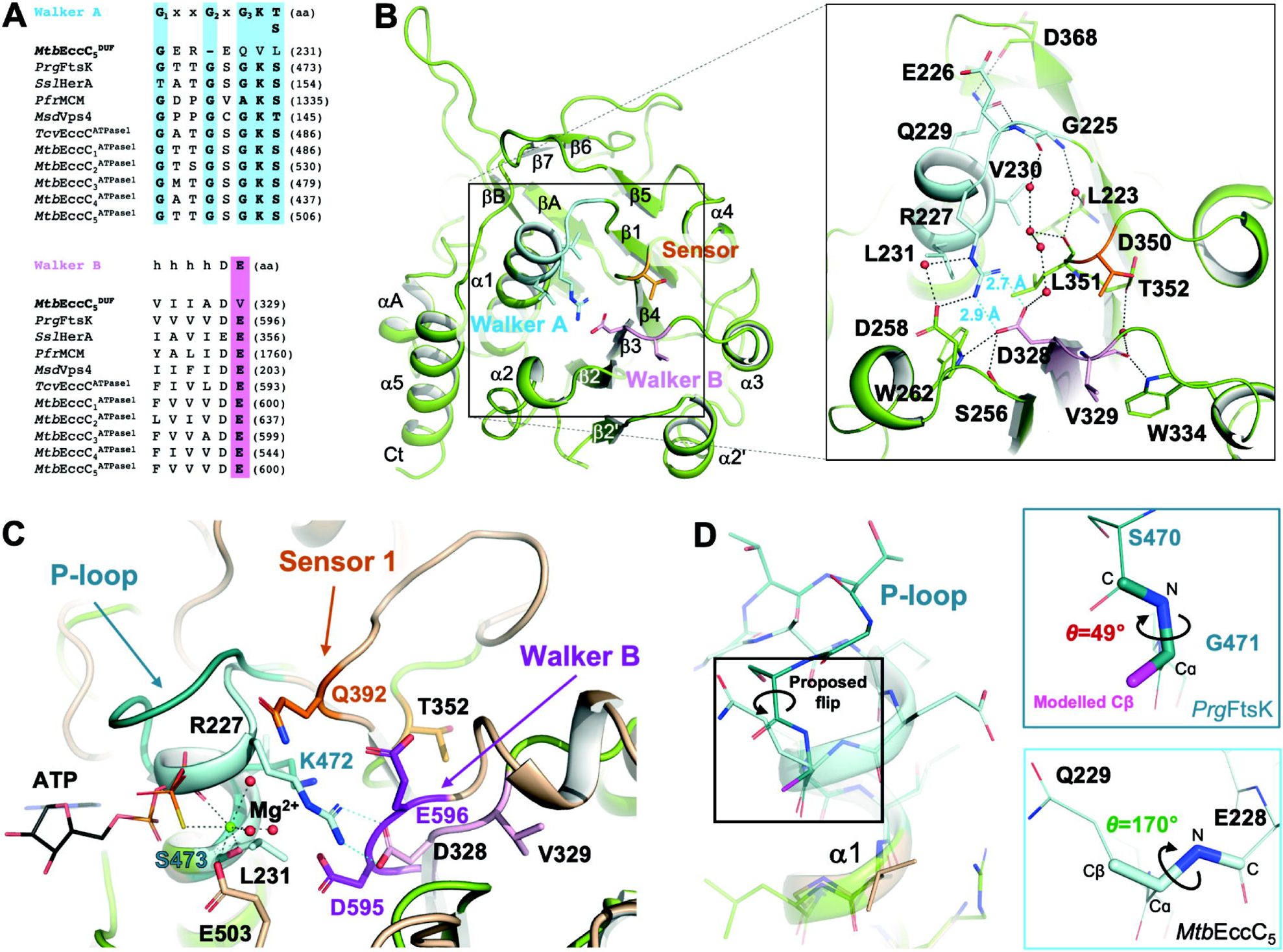
(A) Sequence alignment of Walker A and B motives of *Mtb*EccC_5_^DUF^ with the canonical ATPase domain of *Prg*FtsK*, Ssl*HerA, *Pfr*MCM, *Msd*Vps4, and *Mtb*EccC_1-5_^D1^, showing the consensus sequence at the top. Key amino acids involved in nucleotide interaction/hydrolysis are highlighted in pink and blue, respectively. (B) A general view of the *Mtb*EccC_5_^DUF^ crystal structure represented as green cartoons is shown (left), as well as details of the putative ATP-binding site (right). Salt-bridge and H-bond interactions are indicated as dashed lines in blue and black, respectively, as well as the bond distances measured between R227 and D328, and E226 and Q229. Walker A, B and Sensor 1 motives are coloured in pink, blue and orange respectively, showing key residues (sticks) and water molecules (red spheres). (C) Superimposition of *Mtb*EccC_5_^DUF^ and *Prg*FtsK (PDB entry 4R7Y, in beige) structures represented as cartoons. Walker A, B and Sensor 1 motives are coloured in light (*Mtb*EccC_5_^DUF^) and dark (*Prg*FtsK) blue, pink and orange, respectively. In *Prg*FtsK, magnesium cation, ATP, and residues key for the interaction with the nucleotide are represented as a purple sphere, lines, and sticks, respectively. In The R227-D328 salt bridge observed in *Mtb*EccC_5_^DUF^, and H-bonds involved in ATP-Mg^2+^ stabilization in *Prg*FtsK are represented as dashed lines in black and blue, respectively. (D) Detail of the Walker A motif in *Mtb*EccC_5_^DUF^ and *Prg*FtsK, showing the favourable dihedral angle C_i_-N_i+1_-Cɑ_i+1_-Cβ_i+1_ observed between E228 and Q229 in the former, versus the non-favourable dihedral angle generated between S470 and G471 of *Prg*FtsK when the glycine is substituted by an alanine residue.

The proposed ATP-binding site should localise at the N-terminus of the ɑ1 helix and the C-terminus of the β3 strand, expected to allocate the P-loop and the Walker B motif, respectively. Sequence analysis of the Walker A shows a highly degenerated motif (‘GEREQVL^231^’) exhibiting a 1-aa shorter sequence missing the conserved G_2_ and G_3_ glycines, and lacking the Lys and Thr/Ser residues key for ATP/Mg^2+^ stabilisation (V230 and L231 in *Mtb*EccC_5_^DUF^) (Fig. 2B). Inspection of *Mtb*EccC_5_^DUF^ crystal model shows an ATP-devoid structure where the degenerated Walker A motif folds as an extended ɑ1 helix instead of the standard P-loop conformation. In this arrangement, ɑ1 would clash with the nucleotide molecule, as reveals the structural superimposition of *Mtb*EccC_5_^DUF^ with FtsK from *P. aeruginosa* (*Prg*FtsK), a close-homologue canonical ATPase. To understand the structural basis causing this extended ɑ1 structure, we investigated the impact of G_2_ and G_3_ variations in *Mtb*EccC_5_^DUF^, as invariant flexible features involved both turns of the P-loop motive. To do so, we modelled *in silico* the replacement of G_3_ by alanine in *Prg*FtsK (G471A), which shows how the newly added Cβ gives place to an unfavoured dihedral angle C_i_-N_i+1_-Cɑ_i+1_-Cβ_i+1_ (*θ* = 49°) between S470 and A471 residues (Fig. 2D). This is supported by the poor occurrence of this torsion angle when searched in the CCDC database (Fig. 2SM). This disfavoured conformation is prevented in *Mtb*EccC_5_^DUF^ by a flip of the E228-Q229 peptide bond leading to the prolonged alpha-helical structure observed in ɑ1. Analogously, the absence of G_2_ should foster an alpha-helical conformation to avoid non-favourable dihedral angles between the two adjacent non-glycine amino acids. Altogether, this points to G_2_ and G_3_ variations as structural factors responsible for the extended ɑ1 helix observed in *Mtb*EccC_5_^DUF^. Inspection of the Walker B motif also shows conformational deviations in *Mtb*EccC_5_^DUF^, which includes a shorter β3-ɑ3 loop and a hydrophobic residue (V329) replacing the catalytic base (Fig. 2B). V329 rests partially buried between β3-ɑ3 and β4-ɑ4 loops in a conformation well stabilised by an H-bond network with nearby residues. This drags the Walker B away from the P-loop by > 2 Å compared to *Prg*FtsK, thus providing the room necessary to allocate the extended ɑ1. This non-canonical arrangement of both Walker motives is further stabilized by numerous direct and water-mediated interactions. These include an H-bond between the side-chain amide group of Q229 and the main-chain nitrogen of E226, and a salt-bridge formed between R227 and D328 side chains, stabilising the extended ɑ1 structure, which are well defined in the electron density maps (Fig. 2SM). Moreover, the short bond distances measured for the salt-bridge interaction indicate this is a strong contact that, altogether, should further challenge structural rearrangements of this non-productive conformation observed in ɑ1 for ATP binding. Superimposition of crystal and cryo-EM structures of *Mtb*EccC_5_ shows overall conservation of the non-canonical arrangement observed in the Walker motives in both models (Fig. 2SM). However, significant differences were found in structural features shaping the architecture of the degenerated ATP-binding site, supported by the electron density and cryo-EM maps (Fig. 2SM). These comprise many direct or water-mediated H-bonds stabilising Walker and Sensor 1 motives, including the interaction between Q229 and E226 that is too long to form an H-bond in *Mtb*EccC_5_^CRYO^; or the water-mediated contact established between R227 and D258 side chains, which seems to assist R227 side chain to adopt an optimal orientation to interact with D328 and was not observed in the cryo-EM model. Moreover, contrasting with *Mtb*EccC_5_^CRYO^, the crystal structure of *Mtb*EccC_5_^DUF^ shows a bidentate salt-bridge between R227 and D328 side chains well defined on the electron density maps that contributes to optimise the interactions stabilising the non-canonical configuration of the Walker A motif (Fig. 2SM). Thus, our structure provides an unambiguous model defining the non-canonical ATP-binding site of *Mtb*EccC_5_ at high resolution, which has allowed us to acquire a detailed description and understanding of key structural features contributing to its degenerated arrangement.

We next analysed the regions expected to allocate the *trans*-acting elements in *Mtb*EccC_5_^DUF^ (Fig. 2SM). No basic residues were observed at the end of ɑ4 as candidates to act as Arg-finger. Only R362 was found in the vicinity and further down the ɑ4 sequence. However, its quite buried position in the structure makes it a poor candidate for functioning as an Arg finger. The helical C-terminal lid, habitually found in prokaryotic AAA + clades, that should allocate Sensor 2 is missing in *Mtb*EccC_5_^DUF^. Besides, inspection of the ɑ3 helix shows no basic residues in this region as potential Sensor 3. Taking all these observations together, our structural analysis of *Mtb*EccC_5_^DUF^ shows the degeneration and absence of *cis*- and *trans*-acting elements in *Mtb*EccC_5_^DUF^, respectively, which would support the lack of ATPase activity in this domain. These observations are in line with our failed attempts to co-crystallise *Mtb*EccC_5_^DUF^ with ATP/Mg^2+^, and structures devoid of nucleotide and magnesium obtained instead.

### 4.2 ATP-binding and ATP-hydrolysis analysis

The structural arrangement of the putative ATP-binding site of *Mtb*EccC_5_^DUF^ resembles the nucleotide-binding pocket observed in the structure of the *Srs*EssC^D3^ domain (PDB entry 6TV1), reported to bind ATP with an affinity in the low millimolar range (Mietrach *et al*., 2020) (Fig. 3A). This poor but detectable binding points to potential structural rearrangements of the ATP-binding site of *Srs*EssC^D3^ enabling the interaction with the nucleotide. Therefore, we analysed the interaction of *Mtb*EccC_5_^DUF^ with ATP/Mg^2+^ in order to explore a possible reorganization of the ATP-binding site that might enable nucleotide binding and hydrolysis in solution using two different biophysical approaches.

**Figure 3.**
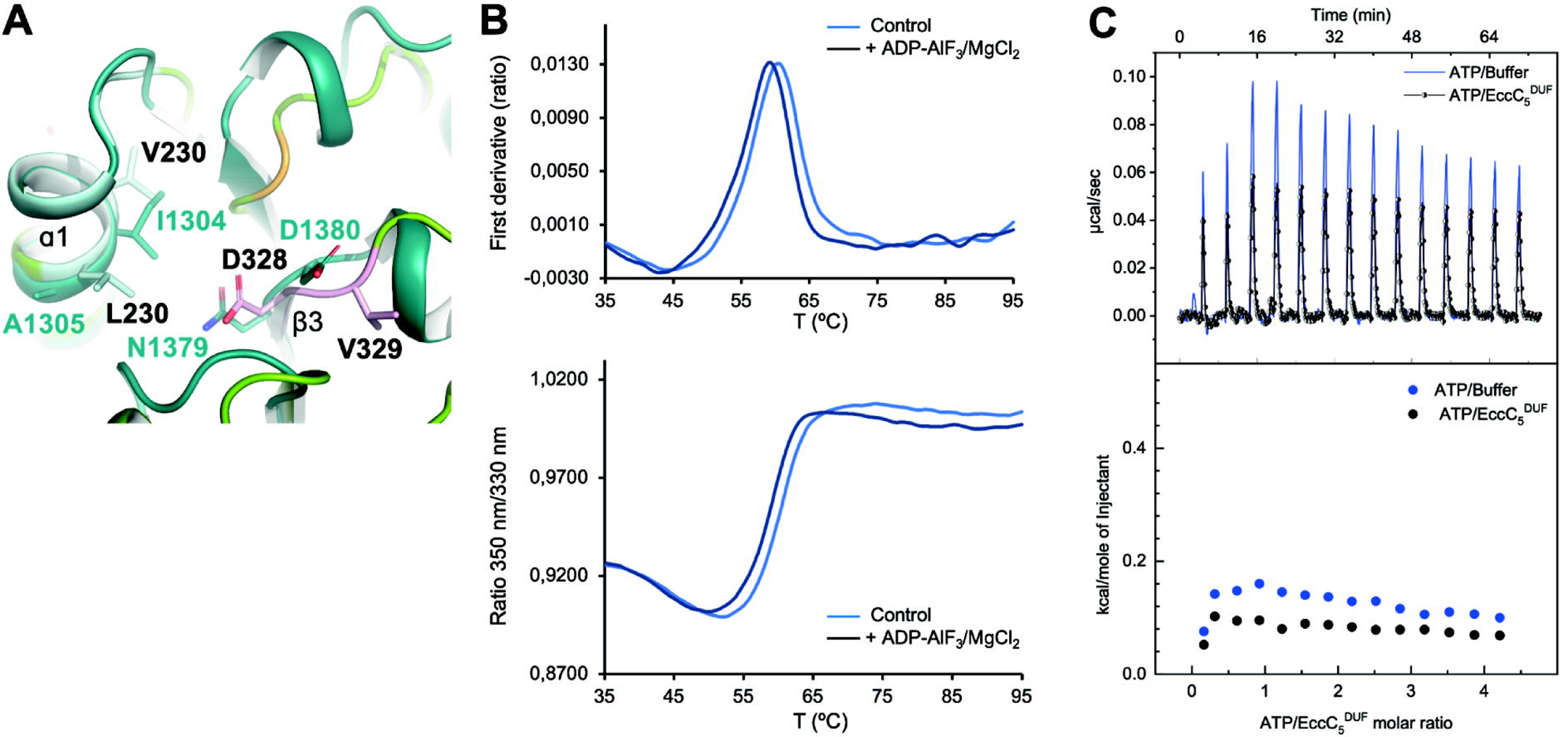
(A) Superimposition of *Mtb*EccC_5_^DUF^ (in green) and *Srs*EssC^D3^ (PDB entry 6VT1, in dark blue) crystal structures showing a detail of their degenerated ATP-binding sites. Residues occupying key positions for nucleotide/magnesium interaction and hydrolysis are labelled and represented as sticks. Walker A, B and Sensor 1 motives of *Mtb*EccC_5_^DUF^ are coloured following the same colour code as in Figure 2. (B) Superimposition of *Mtb*EccC_5_^DUF^ DSF profiles in the absence (control) and presence of ADP-AlF_3_ (4 mM) and Mg^2+^ (5 mM). The failure of ADP-AlF_3_/Mg^2+^ to stabilize the domain structure against thermal denaturation was consistent with no nucleotide binding.(C) ITC titration of ATP (1.3 mM) into EccC_5_^DUF^ (60 μM) and buffer C (black and blue data, respectively). Upper and lower panels show raw thermograms, and normalized heat effect per mole of ATP injected *vs.* ATP/EccC_5_^DUF^ molar ratio, respectively.

Firstly, the interaction of *Mtb*EccC_5_^DUF^ with ADP-AlF_3_ (used as ATP analogue) was assessed by DSF in the presence of magnesium. Despite the high ADP-AlF_3_:EccC_5_^DUF^ concentration used (∼ 4mM), the experiments showed that the difference between *T*_i_ values in the presence (59.13 ± 0.04 °C) and in the absence (60.46 ± 0.04 °C) of nucleotide was slightly negative (around −1.3 °C) (Fig. 3B) and very close to the device’s error limit, thus pointing to the absence of binding. Nucleotide interaction and ATP hydrolysis were next assessed by ITC in the presence of magnesium. The heat released upon titration of ATP into *Mtb*EccC_5_^DUF^ was comparable to the thermal effect of ATP dilution in the range of concentrations tested (Figure 3C). Additionally, no thermal power deflection (proportional to the potential substrate concentration in the cell) was observed after each ATP injection, as would be expected if nucleotide was being hydrolysed (Menéndez, 2020). Altogether ITC and DSF results supports the lack of ATPase activity by *Mtb*EccC_5_^DUF^.

### 4.3 Nucleotide-binding site in other EccC^DUF^ domains

Given the structural degeneration observed in the *Mtb*EccC_5_^DUF^ structure, and its observed inability to interact with ATP using X-ray crystallography, DSF and ITC, we decided to examine the putative ATP-binding site in other DUF domains. With this purpose, homologue EccC^DUF^ and orthologue EssC^DUF^ domains from mycobacterial and non-mycobacterial species were used for sequence alignment (Fig. 4A), which showed degeneration of the Walker A motif in all the domains inspected. Analogously to *Mtb*EccC_5_^DUF^, these motifs present a 1-aa shorter sequence, where conserved residues G_2_ and G_3_ are missing and substituted by a non-glycine amino acid, respectively. Also, they lack both the lysine and threonine/serine residues required for ATP/Mg^2+^ interaction. DUF domains inspected also exhibit degeneration of their Walker B motives, where the catalytic glutamate is replaced by other hydrophobic or polar amino acids (Rosenberg *et al*., 2015; Zoltner *et al*., 2016; Wang *et al*., 2020). Interestingly, analysis of *Mxp*EccC_5_^DUF^ and *Msm*EccC_3_^DUF^ structures showed that their Walker motifs adopt a similar arrangement to the one observed in *Mtb*EccC_5_^DUF^, including an extended ɑ1 helix (Fig. 4A). Analogously to *Mtb*EccC_5_^DUF^, the prolonged ɑ1 helix can be explained by the amino acid variations observed at G_2_ and G_3_ positions with respect to canonical ATPases, which should elicit a helical conformation versus the P-loop structure. This structural reorganisation was also found in the structural models predicted for *Mtb*EccC_1_a^DUF^, *Mtb*EccC_2_^DUF^ and *Mtb*EccC_4_^DUF^ and other six DUF domains, all of which present the same extended conformation of ɑ1 (Fig. 4 and 4SM). Inspection of Walker motifs in *Mxp*EccC ^DUF^ and *Msm*EccC_3_^DUF^ structures also reveal an arginine-aspartate salt-bridge connecting ɑ1 with the C-terminus of β3 (Fig. 4B). The salt-bridge observed in *Mxp*EccC_5_^DUF^ involves an arginine at the N-terminus of ɑ1, analogously to *Mtb*EccC_5_^DUF^, while the salt bridge in *Msm*EccC_3_^DUF^ is formed by an arginine substituting the canonical threonine/serine of the Walker A motif (Fig. 4A). Out of the fifteen sequences analysed, five mycobacterial DUF domains exhibit an arginine/lysin and aspartate that could potentially form a salt bridge, also predicted in the structural models of *Mtb*EccC_2_^DUF^ and *Mtb*EccC_4_^DUF^, where it is likely to contribute to stabilise this unproductive arrangement for ATP-Mg^2+^ recognition (Fig. 4B). Thus, our analysis shows that the non-canonical structure observed in *Mtb*EccC_5_^DUF^ Walker motives is present in the cryo-EM structures of *Mxp*EccC_5_^DUF^ and *Msm*EccC_3_^DUF^, as well as in the predicted models available for other DUF of T7Sa and T7Sb systems. Altogether these observations support that DUFs are degenerated ATPase domains exhibiting a non-functional structure for ATP-binding and thus a poor or null ATPase activity.

**Figure 4.**
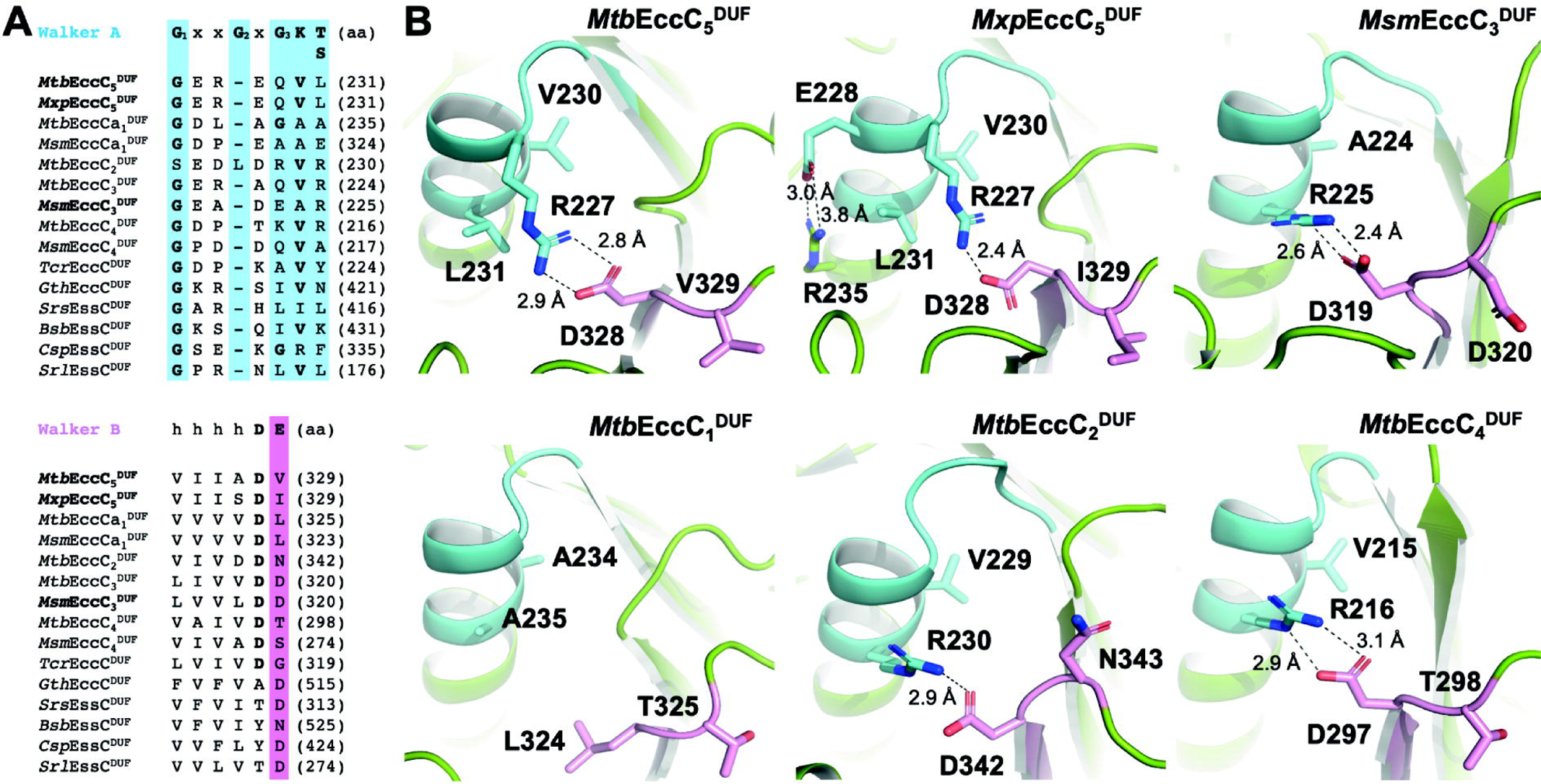
Analysis of putative ATP/Mg^2+^-binding site in EccC^DUF^ domains of ESX1-5 systems from *Mtb*, *Msm* and *Mxp*. (A) Sequence alignment of Walker A and B motives coloured in blue and pink, respectively. (B) Structural details of Walker A and B motives observed in *Mtb*EccC_5_^DUF^ (crystal structure), *Mxp*EccC_5_^DUF^ (PDB entry 7B9S), *Msm*EccC_3_^DUF^ (PDB entry 6GSX), *Mtb*EccC_1_^DUF^ (AlphaFold model, Uniprot entry P9WNB3), *Mtb*EccC_2_^DUF^ (AlphaFold model, Uniprot entry O05450), and *Mtb*EccC_4_^DUF^ (AlphaFold model, Uniprot entry P9WNA7) represented in cartoon and following the same colour code as in (A). Amino acids substituting key residues for ATP-Mg^2+^ interaction, or involved in H-bond/salt-bridge interactions are depicted as sticks.

### 4.4 Hexameric model of *Mtb*EccC_5_ N-terminal region

The lack of ATPase function in EccC^DUF^ domains here proposed raises important questions about their role in the secretion process. The relevance of these questions is particularly stressed by site-directed mutagenesis studies conducted on *Msm*ESX3, where mutations of both aspartate residues of the Walker B motif of *Msm*EccC_3_^DUF^ to alanine (D319A and D320A) were shown to abrogated secretion (Famelis *et al*., 2019). This observation leads us to speculate that these mutations might affect protein:protein interactions that are key for secretion, and would be stablished between the DUF domain and other cytosolic components, or between adjacent DUF domains. Given the conserved ATPase-like fold of DUF, it is plausible to expect that this domain has retained the ability to multimerise, analogously to FtsK homologues and as proposed for EccC^D1-3^ domains as well (Rosenberg *et al*., 2015; Miller & Enemark, 2016). This hypothesis aligns with the size-exclusion chromatography profile observed during the production *Mtb*EccC_5_^DUF^, which indicates the presence of monomeric, dimeric and hexameric species (Fig. 5SM). To evaluate this hypothesis, and based on the multimerised structure of *Prg*FtsK (PDB entry 2IUU), we modelled the structure of the N-terminal region of *Mtb*EccC_5_ in an hexameric state, including TM, stalk and DUF domains, using the crystal coordinates of *Mtb*EccC_5_^DUF^ and *Mtb*EccC_5_^CRYO^ (PDB entry 7NPR) (Fig. 5A). According to this model, *Mtb*EccC_5_^DUF^ domains would form a closed ring, with stalk and TM regions located perpendicular to the DUF domains. Analysis of the interface between adjacent DUFs shows that both Walker A, B and Sensor1 motives are located near the neighbouring monomer, but they do not participate in inter-domain contacts. However, direct interactions via residues at β2-ɑ2 loop, ɑ2 and β5-β6 loop regions were identified. Interestingly, the β2-ɑ2 region locates adjacent to Walker A and B motives, with which it forms direct interactions through several residues that include R227 and D328 (Fig. 5B). Based on these interactions, mutations in R227 and D328 and/or other amino acids of both Walker motives might impact the structure of the β2-ɑ2 region and, by extension, *Mtb*EccC_5_^DUF^ multimerisation. Of note, this would explain as well the lack of secretion observed in both Walker B mutants, D319A and D320A, of *Msm*EccC_3_^DUF^ (Famelis et al., 2019), that correspond to Walker B residues D328 and V329 in *Mtb*EccC_5_, respectively. Regarding TM and stalk regions, they are located perpendicularly, at the edge of the ring formed by *Mtb*EccC_5_^DUF^ domains, with the TM regions of opposite monomers separated by a distance of ∼86 Å. This disposition notably contrasts with EccC_3_ and EccC_5_ organization observed in *Msm*ESX3, *Mtb*ESX5 and *Mxp*ESX5 structures, where the TM regions multimerise to form a helical bundle that connects, via stalk region, with the DUF domains laying away from each other (Famelis *et al*., 2019; Bunduc *et al*., 2021; Beckham *et al*., 2021) (Fig. 5C). Of note, the TM helical bundle locates at the centre of the membrane region where it contributes to a close conformation to the membrane pore. Thus, the aperture of TM and stalk regions depicted by our model suggests a potential conformational change of the N-terminal region of *Mtb*EccC_5_^DUF^ that would explain how the membrane pore opens through the hexamerisation of the DUF domain (Fig. 5C). Altogether, our *Mtb*EccC_5_ model suggests that DUF hexamerisation may play a critical role in the aperture of the membrane pore. This multimerisation would be triggered by the interplay of DUFs with EccC^D1-3^ ATPase domains, thus linking ATP hydrolysis and effector recognition with the opening of the membrane pore necessary for secretion. Certainly, the function here proposed for DUF domains will request further studies to be verified, and raises additional outstanding questions that require our attention, such as how ATPase hydrolysis, effector recognition and multimerization of D1-3 ATPase domains are coupled with the proposed DUF hexamerisation and the aperture of the membrane pore.

**Figure 5.**
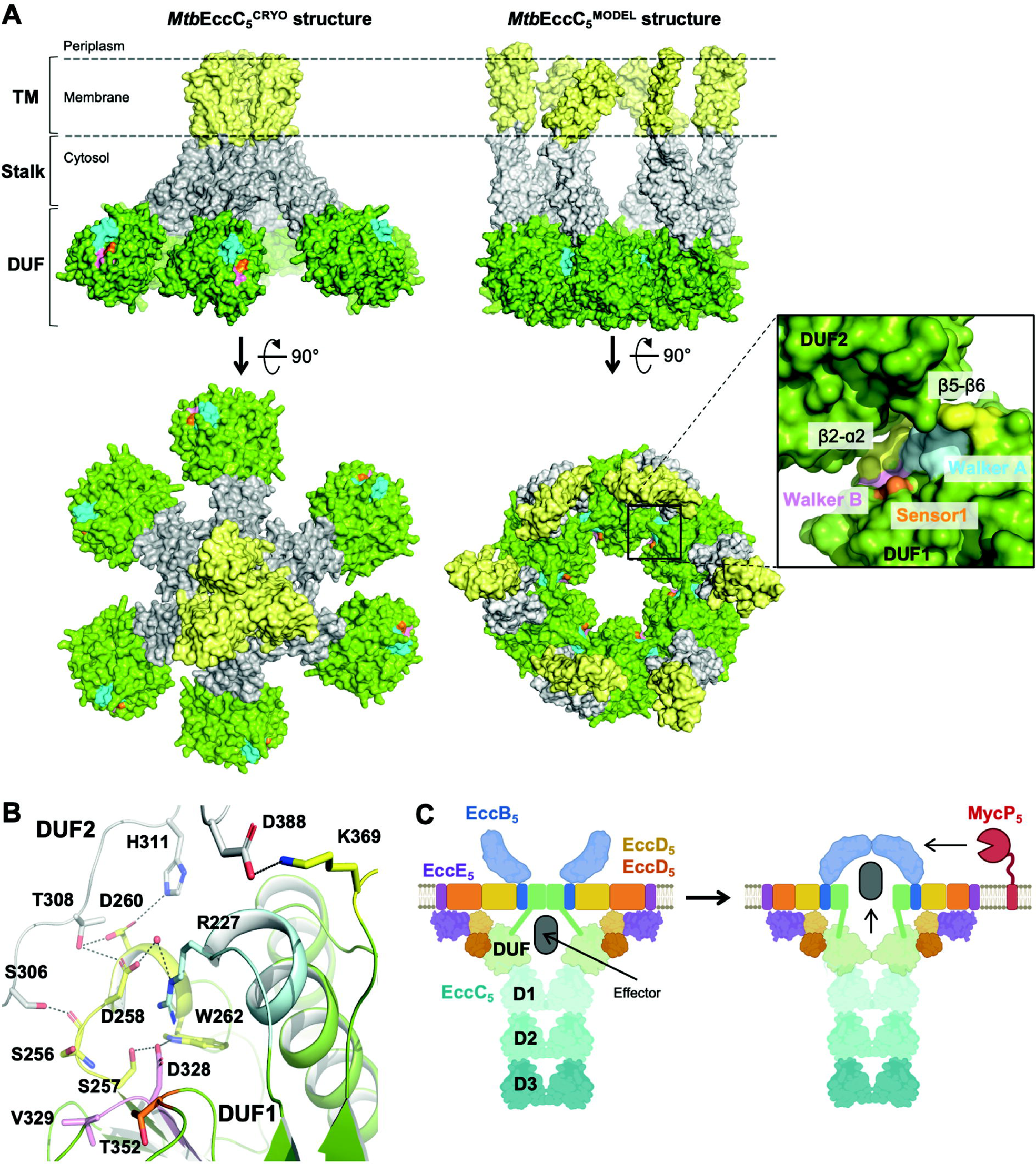
(A) Structural comparison of EccC_5_ arrangement observed in *Mtb*ESX5 structure (PDB entry 7NPR) with our model of *Mtb*EccC_5_ N-terminal region multimerising as a hexamer. A surface representation of *Mtb*EccC_5_ molecules is shown, with transmembrane, stalk and DUF domains coloured in yellow, grey and green, respectively. Walker A, B and Sensor1 motives are coloured in cyan, pink, and orange respectively. A zoom view showing the molecular interface between adjacent DUF monomers in our model is shown at the right box, where identified contact regions have been highlighted in yellow. (B) Details of the interface between DUF1 (grey) and DUF2 (colour code as in A) adjacent molecules is shown, with residues establishing contacts between both monomers and with Walker A and B motives represented as sticks, and their corresponding interactions are indicated as dashed lines. Water molecules are represented as red spheres. (C) Model showing the aperture of the TM region of *Mtb*EccC_5_ associated to the proposed multimerization of *Mtb*EccC_5_^DUF^, where this domain would transit from a non-multimerised form to an hexameric state to enable the opening of the membrane pore.

## 5 Conclusions

Here we report the crystallographic structure of the *Mtb*EccC_5_^DUF^ domain at high resolution, which reveals an ATP/Mg^2+^-free structure with both high degeneration and absence of *cis* and *trans*-acting motives required for ATP-binding and hydrolysis, respectively. Among the most remarkable features, we find the absence of the catalytic glutamate required for the nucleophilic attack over the ATP, and a non-canonical fold of the Walker A motif. Instead of the typical P-loop conformation, the Walker A motif adopts an extended alpha-helical arrangement that would clash with the nucleotide. Although overall conservation of this arrangement is also observed in *Mtb*EccC_5_^CRYO^, our crystal structure provides unseen and unambiguously defined interactions shaping the non-canonical ATP-binding site observed. Moreover, our structural analysis of *Mtb*EccC_5_^DUF^ has led us to identify the structural features underlying this arrangement. They include the variations of G_2_ and G_3_ observed in the Walker A motif of *Mtb*EccC_5_^DUF^, which introduce geometrical restraints that favour the extended ɑ1 helical structure instead of the canonical P-loop conformation. We also validate the lack of interaction of *Mtb*EccC_5_^DUF^ with the nucleotide in solution using DSF and ITC, which aligns with our futile attempts to co-crystallise this domain with ATP-Mg^2+^ that further supports the inability of the DUF domain to bind ATP. Our study also show that the degenerated ATP-binding site observed in *Mtb*EccC_5_^DUF^ is present in the experimental and predicted structures of other homologue domains from T7Sa and T7Sb systems, which supports the notion that DUFs are degenerated ATPase domains, non-functional for nucleotide interaction/hydrolysis. DUF domains emerge, however, as pivotal elements in the secretion process, as illustrated by the mutagenesis studies of the Walker B motif of *Msm*EccC_3_^DUF^ leading to abrogation of the *Msm*ESX3 secretion. Putting all these observations together and based on a structural model of *Mtb*EccC_5_ in a hexameric form, we proposed DUF domains as key structural elements involved in the opening of the membrane pore during the secretion process. This hypothesis, which requires of further studies to be validated, also opens new important questions such as how ATPase hydrolysis, effector recognition and multimerization of D1-3 ATPase domains couple with the proposed hexamerisation of DUF domains and the aperture of the membrane pore.

## 6. Acknowledgments

This work was funded by the *Atracción de Talento* (*Modalidad 1*) program from the Regional Government of Madrid (Spain) (reference 2019-T1/BMD-14774). We also thank the Synchrotron Radiation Facility ALBA (Barcelona, Spain) for the time allocated for X-ray data collection under proposal 2022096995, and BL13-XALOC beamline staff for their assistance in data collection. M. Menendez was supported by the CIBER of Respiratory Diseases (CIBERES), an initiative from the Spanish Institute of Health Carlos III (ISCIII). We also want to thank Dr. Lourdes Infantes San Mateo from the IQF-CSIC, for her input in the structural analysis presented in this work.

## Legends. Supplemental material figures

**Figure 2SM.** (A) Structural superimposition of both crystallographic and cryo-EM *Mtb*EccC_5_^DUF^ models coloured in Figure 2 and grey, respectively. In the case of *Mtb*EccC_5_^CRYO^, N-terminal, stalk and TM regions are coloured in red, dark grey and yellow, respectively. A zoomed view showing the structural conservation of both the Walker A and B motives in both models is shown (right), including the salt-bridge R227-D328 and the H-bond interaction E226-Q229 indicated as dashed lines coloured in blue (crystal model) and grey (cryo-EM model). (B) Mogul search results obtained for torsion angles C_S470_-N_A471_-Cɑ _A471_-Cβ _A471_ of *Prg*FtsK, when G471 is substituted by an alanine, and C_E228_-N_Q229_-Cɑ _Q229_-Cβ _Q229_ in *Mtb*EccC_5_^DUF^ crystal structure. (C) and (D) Detail of both Walker A and B motives in the *Mtb*EccC_5_^DUF^ crystal structure showing the 2Fo-Fc electron density map (contoured at a sigma value of 1.0). Colour code as in Figure 1. (D) Detail of both Walker A and B motives in *Mtb*EccC_5_^DUF^ model reported by cryo-EM showing the EM map at 3.27 Å resolution (PDB entry 7NPT) (contoured at a level value of 10.0). (E) Cartoon and surface representation *Mtb*EccC_5_^DUF^, showing details of the regions expected to allocate Arg-finger and Sensor 3 motives (highlighted in orange).

**Figure 4SM.** (A) Structural superimposition showing a detail of the Walker A and B motives predicted for *Mtb*EccC_5_^DUF^, *Tcr*EccC^DUF^*, Gth*EccC^DUF^*, Srs*EssC^DUF^*, Bsb*EssC^DUF^*, Csp*EssC^DUF^,and *Srl*EssC^DUF^. (B) Size-exclusion chromatography profile obtained during the purification of the *Mtb*EccC_5_^DUF^ construct used in this study.

**Figure 5SM.** (A) Size-exclusion chromatography profile obtained during the purification of the *Mtb*EccC_5_^DUF^ construct used in this study.

## Bibliography

Adams, P. D., Afonine, P. V., Bunkóczi, G., Chen, V. B., Davis, I. W., Echols, N., Headd, J. J., Hung, L. W., Kapral, G. J., Grosse-Kunstleve, R. W., McCoy, A. J., Moriarty, N. W., Oeffner, R., Read, R. J., Richardson, D. C., Richardson, J. S., Terwilliger, T. C. & Zwart, P. H. (2010). Acta Cryst D 66, 213–221.

Beckham, K. S. H., Ritter, C., Chojnowski, G., Ziemianowicz, D. S., Mullapudi, E., Rettel, M., Savitski, M. M., Mortensen, S. A., Kosinski, J. & Wilmanns, M. (2021). Sci Adv 7, eabg9923.

Bricogne G., Blanc E., Brandl M., Flensburg C., Keller P., Paciorek W., Roversi P, Sharff A., Smart O.S., Vonrhein C. & Womack T.O. (2017). BUSTER v.2.11.2, https://www.globalphasing.com/.

Bunduc, C. M., Bitter, W. & Houben, E. N. G. (2020). Annu Rev Microbiol 74, 315–335.

Bunduc, C. M., Fahrenkamp, D., Wald, J., Ummels, R., Bitter, W., Houben, E. N. G. & Marlovits, T. C. (2021). Nature 593, 445–448.

Caillat, C., Macheboeuf, P., Wu, Y., McCarthy, A. A., Boeri-Erba, E., Effantin, G., Göttlinger, H. G., Weissenhorn, W. & Renesto, P. (2015). Nat Commun 6, 8781.

Chen, V. B., Arendall, W. B., Headd, J. J., Keedy, D. A., Immormino, R. M., Kapral, G. J., Murray, L. W., Richardson, J. S. & Richardson, D. C. (2009). Acta Cryst D 66, 12–21.

Crosskey, T. D., Beckham, K. S. H. & Wilmanns, M. (2020). Prog Biophys Mol Biol 152, 25–34.

Emsley, P., Lohkamp, B., Scott, W. G. & Cowtan, K. (2010). Acta Cryst D 66, 486–501.

Evans, P. R. & Murshudov, G. N. (2013). Acta Cryst D 69, 1204–1214.

Famelis, N., Geibel, S. & Van Tol, D. (2023). Biol Chem 404, 691–702.

Famelis, N., Rivera-Calzada, A., Degliesposti, G., Wingender, M., Mietrach, N., Skehel, J. M., Fernandez-Leiro, R., Böttcher, B., Schlosser, A., Llorca, O. & Geibel, S. (2019). Nature 576, 321–325.

Houben, E. N. G., Korotkov, K. V. & Bitter, W. (2014). Biochimica et Biophysica Acta 1843, 1707–1716.

Jumper, J., Evans, R., Pritzel, A., Green, T., Figurnov, M., Ronneberger, O., Tunyasuvunakool, K., Bates, R., Žídek, A., Potapenko, A., Bridgland, A., Meyer, C., Kohl, S. A. A., Ballard, A. J., Cowie, A., Romera-Paredes, B., Nikolov, S., Jain, R., Adler, J., Back, T., Petersen, S., Reiman, D., Clancy, E., Zielinski, M., Steinegger, M., Pacholska, M., Berghammer, T., Bodenstein, S., Silver, D., Vinyals, O., Senior, A. W., Kavukcuoglu, K., Kohli, P. & Hassabis, D. (2021). Nature 596, 583–589.

Kabsch, W. (2010). Acta Cryst D 66, 125–132.

Leipe, D. D., Wolf, Y. I., Koonin, E. V. & Aravind, L. (2002). J Mol Biol 317, 41–72.

Li, N., Zhai, Y., Zhang, Y., Li, W., Yang, M., Lei, J., Tye, B. K. & Gao, N. (2015). Nature 524, 186–191.

Massey, T. H., Mercogliano, C. P., Yates, J., Sherratt, D. J. & Löwe, J. (2006). Mol Cell 23, 457–469.

McCoy, A. J., Grosse-Kunstleve, R. W., Adams, P. D., Winn, M. D., Storoni, L. C. & Read, R. J. (2007). J Appl Crystallogr 40, 658–674.

Menéndez, M. (2020). Encyclopedia of Life Sciences 1, 113–127.

Mietrach, N., Damián-Aparicio, D., Mielich-Süss, B., Lopez, D. & Geibel, S. (2020). J Bacteriol 202, 646–665.

Miller, J. M., Arachea, B. T., Epling, L. B. & Enemark, E. J. (2014). Elife 3, e03433.

Miller, J. M. & Enemark, E. J. (2016). Archaea 2016, ID 9294307.

Ogura, T., Whiteheart, S. W. & Wilkinson, A. J. (2004). Journal of Structural Biology 146, 106–112.

Poweleit, N., Czudnochowski, N., Nakagawa, R., Trinidad, D., Murphy, K. C., Sassetti, C. & Rosenberg, O. S. (2019). Elife 8, e52983.

Rosenberg, O. S., Dovala, D., Li, X., Connolly, L., Bendebury, A., Finer-Moore, J., Holton, J., Cheng, Y., Stroud, R. M. & Cox, J. S. (2015). Cell 161, 501–512.

Rzechorzek, N. J., Blackwood, J. K., Bray, S. M., Maman, J. D., Pellegrini, L. & Robinson, N. P. (2014). Nat Commun 5, 5506.

Schmidt, H., Gleave, E. S. & Carter, A. P. (2012). Nature Structural & Molecular Biology 19, 492–497.

Sievers, F., Wilm, A., Dineen, D., Gibson, T. J., Karplus, K., Li, W., Lopez, R., McWilliam, H., Remmert, M., Söding, J., Thompson, J. D. & Higgins, D. G. (2011). Mol Syst Biol 7, 539.

Wang, S., Zhou, K., Yang, X., Zhang, B., Zhao, Y., Xiao, Y., Yang, X., Yang, H., Guddat, L. W., Li, J. & Rao, Z. (2020). Protein Cell 11, 124–137.

World Health Organization (2022). Global Tuberculosis Report 2020, https://www.who.int/teams/global-tuberculosis-programme/tb-reports/global-tuberculosis-report-2022.

Zoltner, M., Ng, W. M. A. V., Money, J. J., Fyfe, P. K., Kneuper, H., Palmer, T. & Hunter, W. N. (2016). Biochem J 473, 1941–1952.

