## Supplementary material for "New insights into the domain of unknown function DUF of EccC_5_, the pivotal ATPase providing the secretion driving force to the ESX5 secretion system": SM text

1. Groups of Crystallography and Structural Biology. Institute of Physical-Chemistry Blas Cabrera, Spanish Research Council, Serrano 119, 28006 – Madrid (Spain)

2. Group of Structure and Thermodynamics of Macromolecules. Institute of Physical-Chemistry Blas Cabrera, Spanish Research Council, Serrano 119, 28006 – Madrid (Spain)

3. Ciber of Respiratory Diseases, ISCIII, Sinesio Delgado 10, 28029 – Madrid (Spain)

*Corresponding author

**Supplemental material legends**

**Figure 2SM**. (A) Structural superimposition of both crystallographic and cryo-EM *Mtb*EccC_5_^DUF^ models coloured in Figure 2 and grey, respectively. In the case of *Mtb*EccC_5_^CRYO^, N-terminal, stalk and TM regions are coloured in red, dark grey and yellow, respectively. A zoomed view showing the structural conservation of both the Walker A and B motives in both models is shown (right), including the salt-bridge R227-D328 and the H-bond interaction E226-Q229 indicated as dashed lines coloured in blue (crystal model) and grey (cryo-EM model). (B) Mogul search results obtained for torsion angles C_S470_-N_A471_-Cɑ _A471_-Cβ _A471_ of *Prg*FtsK, when G471 is substituted by an alanine, and C_E228_-N_Q229_-Cɑ _Q229_-Cβ _Q229_ in *Mtb*EccC_5_^DUF^ crystal structure. (C) and (D) Detail of both Walker A and B motives in the *Mtb*EccC_5_^DUF^ crystal structure showing the 2Fo-Fc electron density map (contoured at a sigma value of 1.0). Colour code as in Figure 1. (D) Detail of both Walker A and B motives in *Mtb*EccC_5_^DUF^ model reported by cryo-EM showing the EM map at 3.27 Å resolution (PDB entry 7NPT) (contoured at a level value of 10.0). (E) Cartoon and surface representation *Mtb*EccC_5_^DUF^, showing details of the regions expected to allocate Arg-finger and Sensor 3 motives (highlighted in orange).

**Figure 4SM**. (A) Structural superimposition showing a detail of the Walker A and B motives predicted for *Mtb*EccC_5_^DUF^, *Tcr*EccC^DUF^*, Gth*EccC^DUF^*, Srs*EssC^DUF^*, Bsb*EssC^DUF^*, Csp*EssC^DUF^,and *Srl*EssC^DUF^. (B) Size-exclusion chromatography profile obtained during the purification of the *Mtb*EccC_5_^DUF^ construct used in this study.

**Figure 5SM**. (A) Size-exclusion chromatography profile obtained during the purification of the *Mtb*EccC_5_^DUF^ construct used in this study.
