## Supplementary figures and images for "New insights into the domain of unknown function DUF of EccC_5_, the pivotal ATPase providing the secretion driving force to the ESX5 secretion system"

### Figure2SM

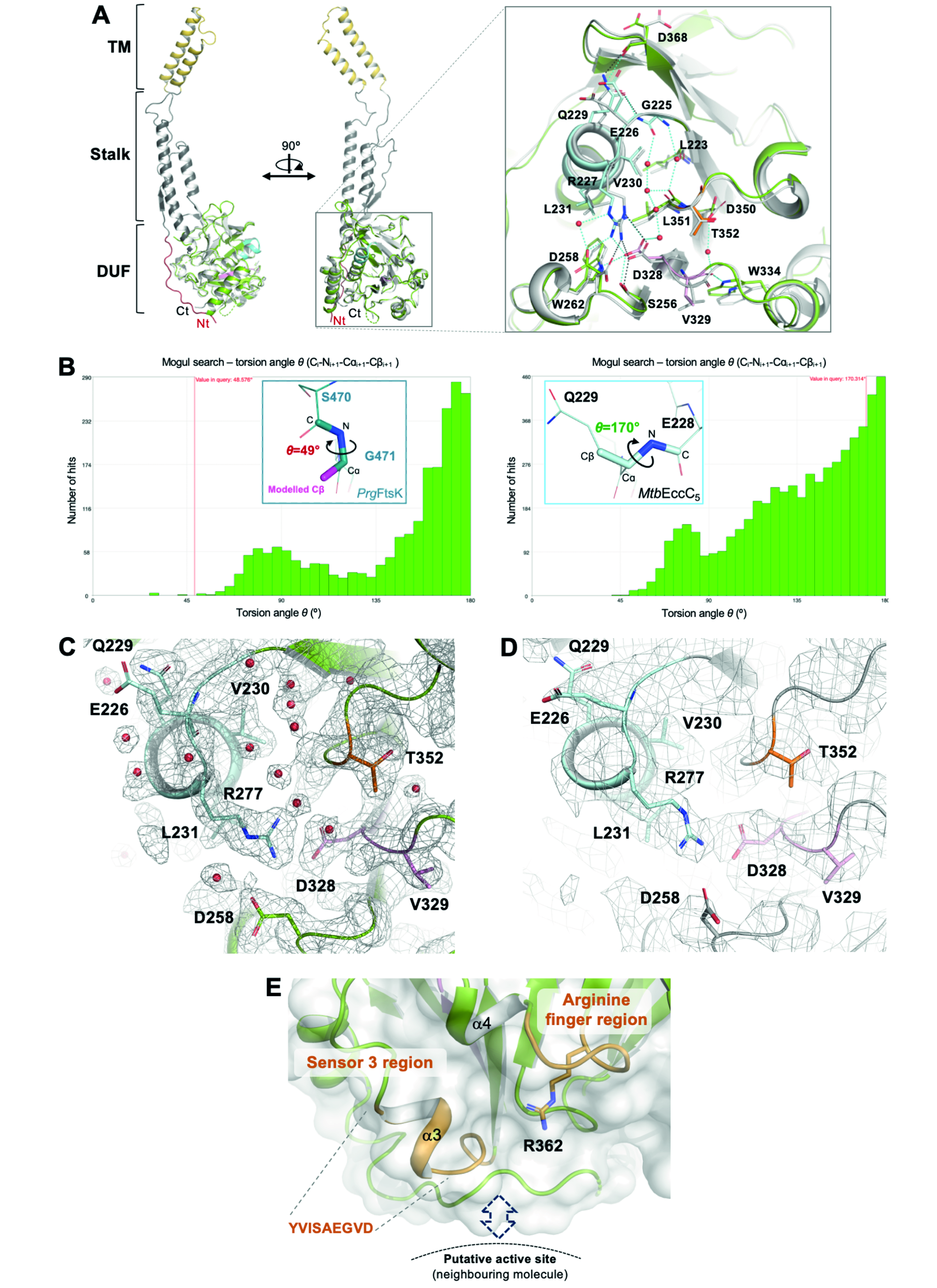

### Figure4SM

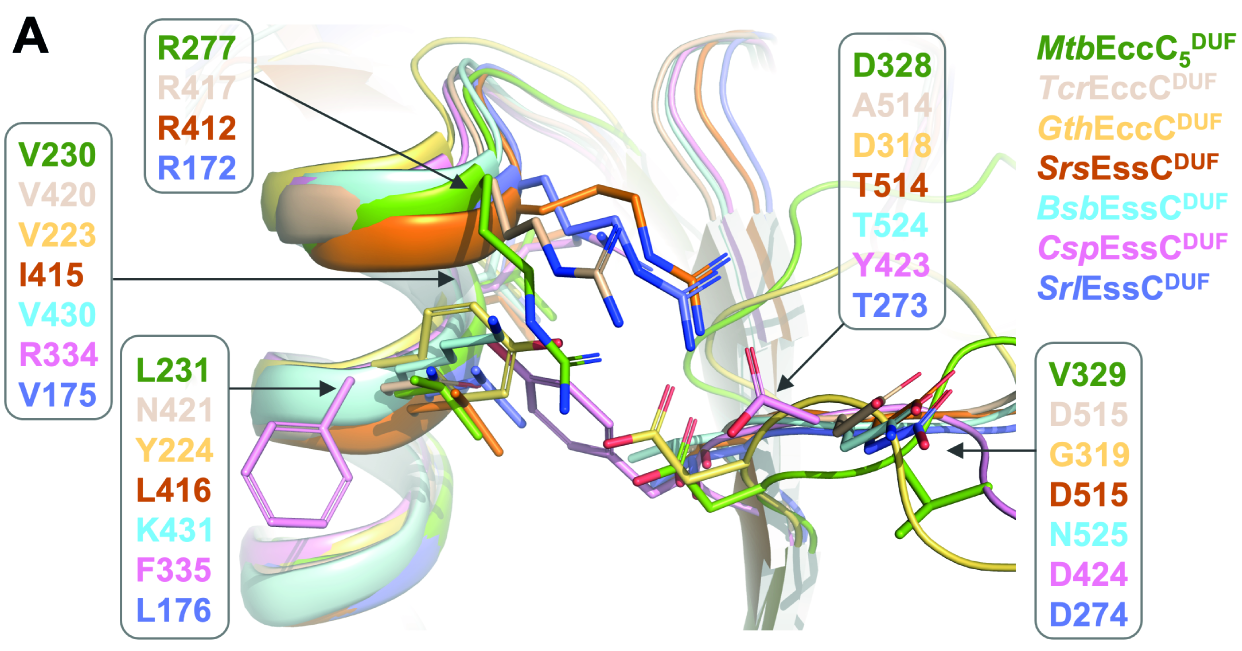

### Figure5SM

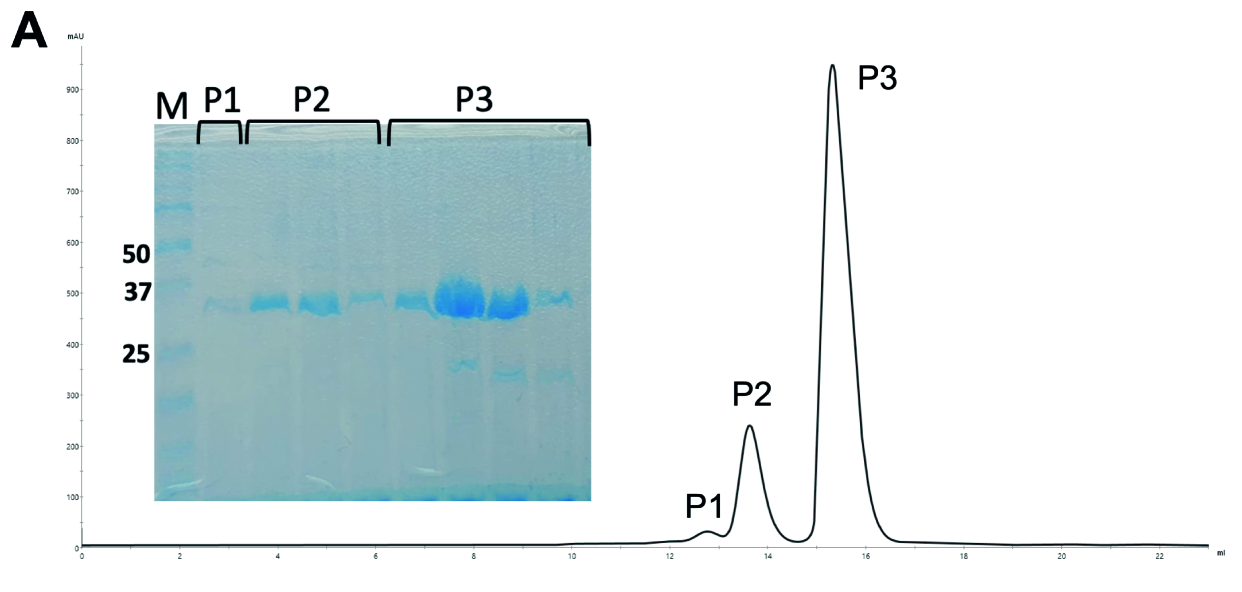
